# Unpacking bedaquiline hetero-resistance: the importance of intermediate profiles for phenotypic drug-susceptibility testing

**DOI:** 10.1101/2025.03.09.642295

**Authors:** Nabila Ismail, Frik Sirgel, Shaheed Vally Omar, Shatha Omar, Marianna de Kock, Claudia Spies, Megan Folkerts, Grant Theron, Dave Engelthaler, John Metcalfe, Robin M Warren

**Author notes:** Contributed equally (Nabila Ismail, Frik Sirgel). Contributed equally (John Metcalfe, Robin M Warren).

## Abstract

Phenotypic drug susceptibility testing (pDST) remains the gold standard for Mycobacterium tuberculosis complex drug resistance determination. Next generation sequencing technologies can identify heteroresistant populations at low frequencies, but little is known about the impact of heteroresistance on bedaquiline (BDQ) pDST results. We simulated heteroresistance using in vitro generated MmpR5 mutants mixed with the progenitor strain at various percentages (1-20%) and did pDST using MGIT960 culture (1 and 2 µg/mL BDQ concentrations). Targeted Next Generation Sequencing (tNGS) was used to quantify the mutant sub-population in growth control tubes, which were expected to maintain the mutant: wild type proportion throughout the assay. Growth units of these growth control tubes were also comparable with minor differences in time-to-positivity between ratio mixtures. Only when intermediate results were considered could BDQ heteroresistance be detected at frequencies of approximately 1% by pDST at a critical concentration of 1 µg/mL using BACTEC MGIT960 coupled with EpiCenter TBeXiST software. The ability of pDST, a widely available DST technique, to reveal the presence of BDQ-resistant subpopulations at the phenotypic testing stage could improve resistance determination and potentially reduce time to effective treatment.

**Importance:** Multidrug resistant tuberculosis (MDR-TB) is estimated to cause up to 19% of all antimicrobial resistance-attributable deaths worldwide. Further, the success rate for the treatment of drug-resistant TB, in the presence of adherence, is poor at only 68%. The advent of bedaquiline (BDQ) has revolutionized MDR-TB care, but BDQ resistance determination is hampered by several obstacles facing both phenotypic and genotypic testing. Specifically for phenotypic susceptibility testing, BDQ-resistant *Mycobacterium tuberculosis* isolates with variants in *MmpR5*, which may display minimal inhibitory concentration values just below the critical concentration or are present at low frequencies (heteroresistance; the presence of mixed mutant and wild-type populations within a specimen), are typically designated as susceptible. This may lead to prescription of an ineffective regimen and amplification of resistance. The BACTEC MGIT960 platform coupled with EpiCenter TBeXiST software for phenotypic testing, which is currently the only routinely used method of BDQ DST, can be used to derive more information about underlying resistant populations. We demonstrate how this is possible through the consideration of intermediate results (i.e., when growth units in a drug-containing tube reach the threshold for resistance but only after a further week of incubation). These intermediate results, commonly disregarded by TB laboratories, could lead to earlier detection of BDQ resistance. This is especially crucial when the genetic mechanism of resistance is unknown, a variant has not been associated with resistance in the interim, and in cases of heteroresistance.

## Introduction

*Mycobacterium tuberculosis* bedaquiline (BDQ) susceptibility determination faces challenges on both genotypic and phenotypic fronts. This is despite the drug having received FDA approval for treatment of adults with pulmonary multi-drug resistant tuberculosis (MDR-TB) in 2012 (1). Mutations in *MmpR5*, a gene encoding a transcriptional repressor (MmpR5), which downregulates transmembrane pump proteins MmpL5-S5, are the most common resistance causing variants. These variants can be found across the length of the 498-bp gene, with no clear hotspot (2). Furthermore, in the 2023 WHO mutation catalog, loss-of-function (LoF) variants in *MmpR5* are graded as Group 2 variants (i.e., “associated with resistance in the interim”), meaning further data is required to statistically support the association with resistance (3). A caveat to this rule, however, is that simultaneous LoF variants in MmpL5-S5 are epistatic (inhibit, mask or suppress the impact of a LoF variant in *MmpR5*) and result in BDQ hyper susceptibility - a phenomenon detected primarily in the Lima, Peru region at a frequency of 43% in sublineage 4.11 isolates (4).

Of the three WHO-recommended targeted next-generation sequencing assays for determination of drug resistance, only one has met the class-based performance criteria for BDQ (5) but none of the cover the *mmpL5-S5* region (5). Additionally, the sensitivity for the use of tNGS to predict a resistant phenotype is currently only 68% for BDQ (6) - highlighting the continued necessity to rely on phenotypic testing to resolve any inadequacies. In certain instances, variants in *MmpR5* (particularly those which result in incomplete repression of the MmpL5/S5 pump) can confer borderline resistant phenotypes (i.e. minimal inhibitory concentration (MIC) values one dilution above or below the critical concentration) - the effect of which could be missed due to the technical variability of phenotypic testing (7, 8). Aside from the long turnaround time, the endorsed critical concentration for BDQ phenotypic susceptibility testing using the MGIT960 platform is based on limited evidence (7). A composite reference standard would be ideal to overcome the limitations of both phenotypic and genotypic BDQ susceptibility testing and can also be used to evaluate the significance of heteroresistance (7).

The impact of heteroresistance was highlighted in the 2023 WHO mutation catalog, when the inclusion of variants with an allele frequency below 75% increased the combined sensitivity of Group 1 and 2 variants (associated with resistance (interim)) by 10.2% and decreased the specificity by only 0.3% (3). In this same catalog, it was also considered that lowering the threshold of variant detection to 25% to account for lower-level BDQ heteroresistance improves prediction of a resistant phenotype (3). Here, we demonstrate a novel application of the BACTEC MGIT960 coupled with the EpiCenter TBeXiST (TB-eXtended individual drug Susceptibility Testing) software, the reference standard, globally available automated culture system for detecting *Mycobacterium tuberculosis* worldwide, to innovatively analyze phenotypic growth unit (GU) and time-to-positivity (TTP) data for early detection of BDQ resistance.

## Methods

### Mutant and progenitor strains

*In vitro* mutants spontaneously generated with a Luria Delbruck assay, using either an ATCC27294 (fully-susceptible H37Rv) or ATCC35828 (PZA-resistant) progenitor strain (BDQ MGIT960 MIC values of 0.5 and 1 µg/mL) were selected on clofazimine (CFZ)-containing media as previously described (9). CFZ exposure is capable of generating *MmpR5* variants, which result in BDQ cross-resistance (10). Mutants were further purified by selecting single colonies on BBL™ 7H11 agar base media (Becton Dickinson, NJ, USA) containing 0.25, 0.5 or 1 µg/mL of clofazimine (Catalog No: A16462, Adooq Biosciences, CA, USA) and supplemented with 10% (v/v) Middlebrook oleic acid, bovine albumin, sodium chloride, dextrose, and catalase (OADC) Growth Supplement (Becton Dickinson) and 0.5% (v/v) glycerol. Complete mutant characterization is in **Table 1**, an overview of the workflow is in **Figure 1** and detailed methods can be found in the supplementary.

**Figure 1:**
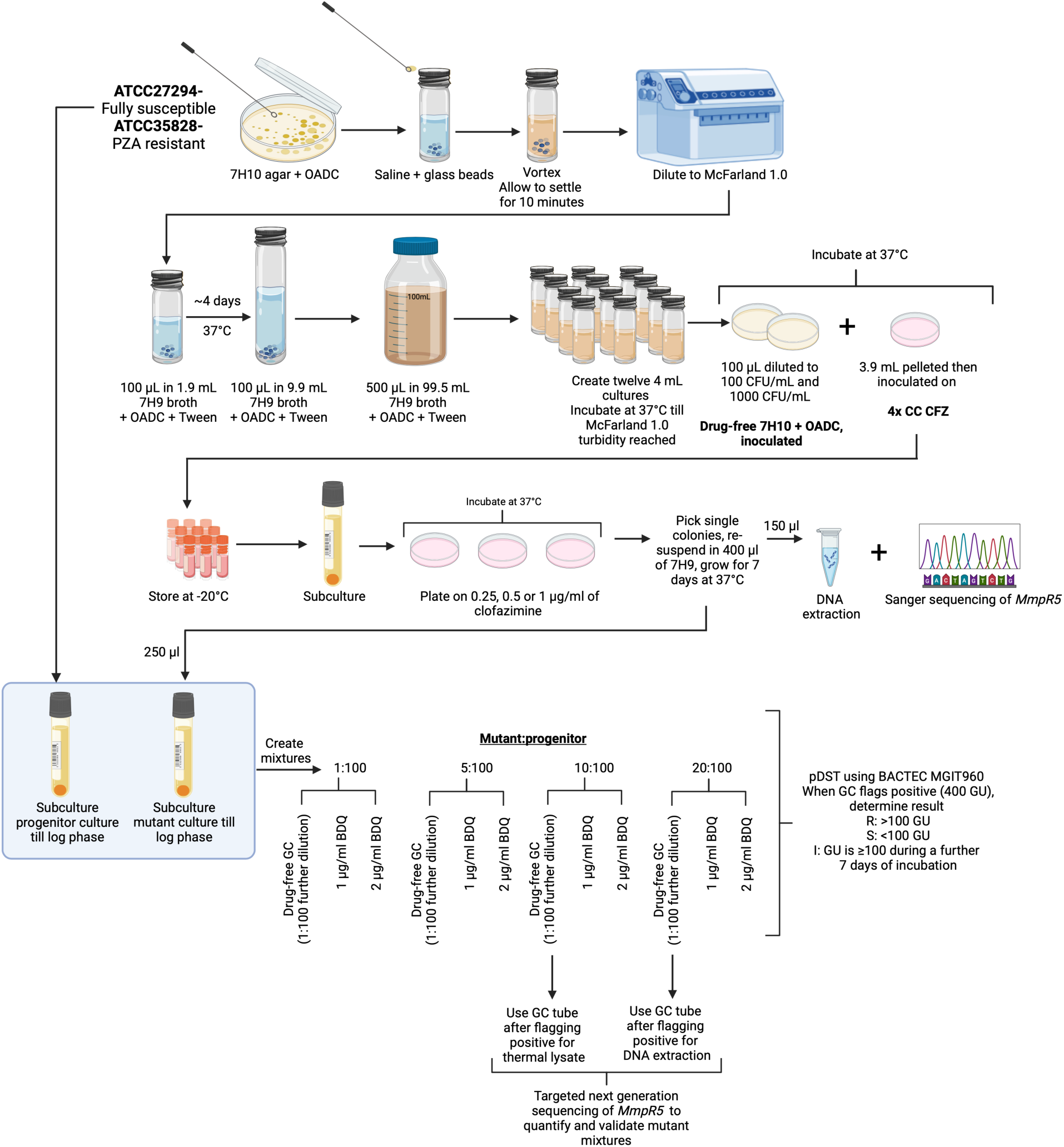
Method flow diagram for creation of clofazimine-resistant *in vitro* mutants, from which single colonies were picked and used for *MmpR5* Sanger sequencing. Pure mutant colonies were then grown to log phase and mixed with progenitor strains, also grown to log phase. Mutant strains are mixed at 1, 5, 10, 20: 100 ratios with progenitor strains. For each mixture, a drug-free control (GC), 1:100 dilution prepared as previously described (12, 13) and drug-containing test tubes (1 and 2 µg/ml BDQ) were set-up. Resistant, susceptible and intermediate pDST results were recorded from the BACTEC960 instrument. Purified DNA and thermal lysates from growth from GC tubes (after flagging positive) for the 10 and 20% mixtures were used for *MmpR5* targeted sequencing to quantify and validate mixtures. Created in https://biorender.com

**Table 1:**
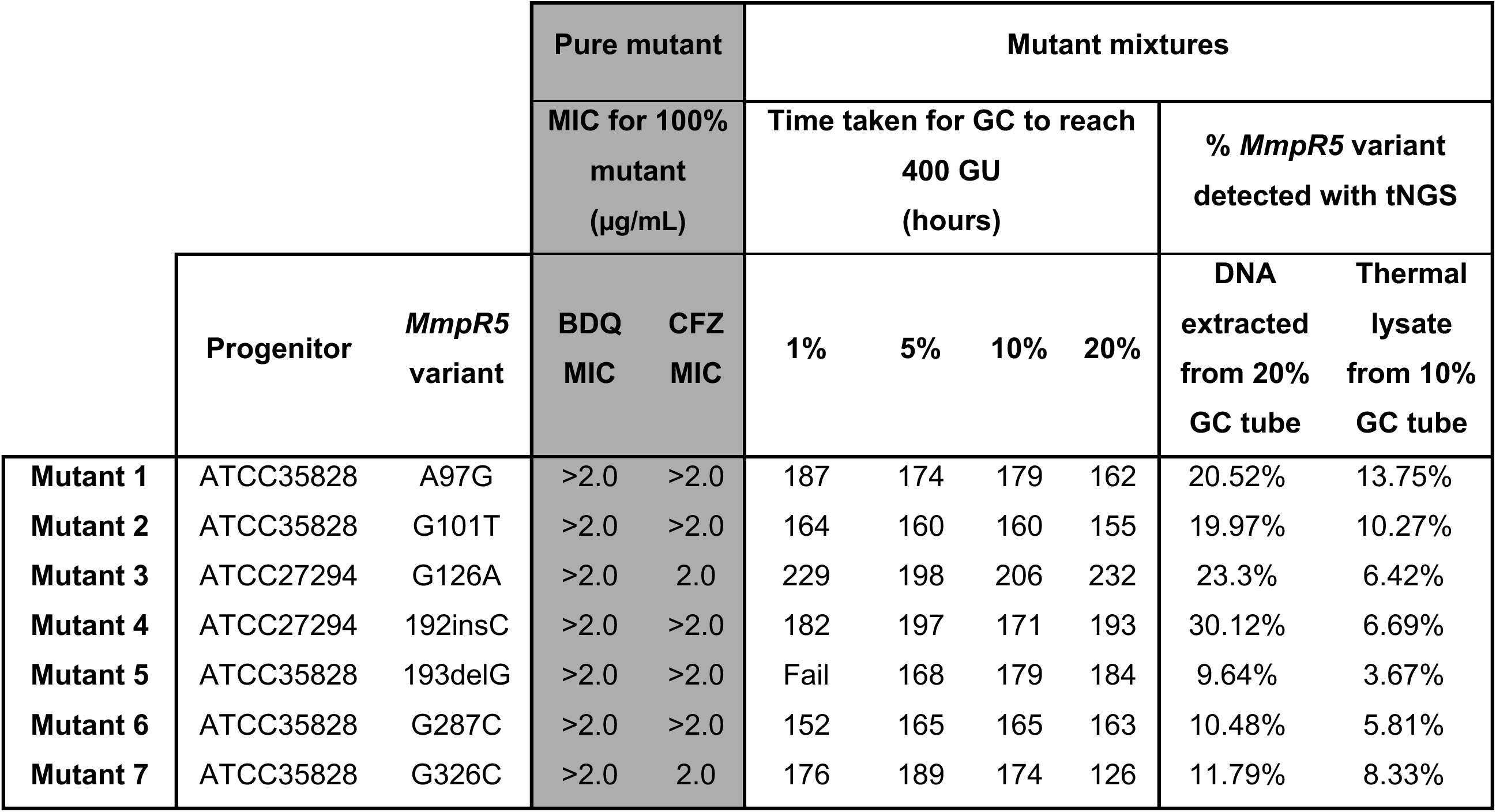
Genotypic and phenotypic data associated with each of the 7 mutants as well as growth units (GU) for growth control (GC) tubes and percentage of mutant DNA from growth control tubes obtained using targeted next generation sequencing (TNGS). Two different progenitor strains were used for *MmpR5* mutant generation; ATCC27294 (BDQ MGIT960 MIC: 0.5 µg/mL) and ATCC35828 (BDQ MGIT960 MIC: 1 µg/mL). These progenitors were used to create corresponding mixtures. Each ture assay comprised of a single GC tube and two drug containing tubes (either 1 or 2 µg/mL BDQ).

|  |  |  | Pure mutant |  | Mutant mixtures |  |  |  |  |  |
| --- | --- | --- | --- | --- | --- | --- | --- | --- | --- | --- |
|  |  |  | MIC for 100%<br>mutant<br>(µg/mL) |  | Time taken for GC to reach<br>400 GU<br>(hours) |  |  |  | % *MmpR5* variant<br>detected with tNGS |  |
|  | Progenitor | *MmpR5*<br>variant | BDQ<br>MIC | CFZ<br>MIC | 1% | 5% | 10% | 20% | DNA<br>extracted<br>from 20%<br>GC tube | Thermal<br>lysate<br>from 10%<br>GC tube |
| **Mutant 1** | ATCC35828 | A97G | >2.0 | >2.0 | 187 | 174 | 179 | 162 | 20.52% | 13.75% |
| **Mutant 2** | ATCC35828 | G101T | >2.0 | >2.0 | 164 | 160 | 160 | 155 | 19.97% | 10.27% |
| **Mutant 3** | ATCC27294 | G126A | >2.0 | 2.0 | 229 | 198 | 206 | 232 | 23.3% | 6.42% |
| **Mutant 4** | ATCC27294 | 192insC | >2.0 | >2.0 | 182 | 197 | 171 | 193 | 30.12% | 6.69% |
| **Mutant 5** | ATCC35828 | 193delG | >2.0 | >2.0 | Fail | 168 | 179 | 184 | 9.64% | 3.67% |
| **Mutant 6** | ATCC35828 | G287C | >2.0 | >2.0 | 152 | 165 | 165 | 163 | 10.48% | 5.81% |
| **Mutant 7** | ATCC35828 | G326C | >2.0 | 2.0 | 176 | 189 | 174 | 126 | 11.79% | 8.33% |

### Sanger sequencing

Each purified colony was inoculated into 400 µL MGIT medium (supplemented with OADC) and grown for 7 days at 37°C. An aliquot of this culture (250 µL) was inoculated into a fresh MGIT tube supplemented with 0.8 ml OADC (MGIT-OADC tube) while the remainder was used as a thermal lysate for PCR amplification of the *M. tuberculosis MmpR5* gene. Successfully amplified PCR products were submitted to the Central Analytical Facility (CAF) of Stellenbosch University for post-PCR purification and Sanger sequencing. Sequencing chromatograms were visually inspected using BioEdit™ Sequence Alignment Editor software version 7.2.6 (11) to characterize *MmpR5* variants selected while ensuring there was no mix of wild-type with mutant sequences-confirming purification of the colonies.

### Drugs

BDQ (Catalog No: A12327, AdooQ Biosciences, CA, USA) and CFZ were formulated in DMSO (Ref: 41639, Sigma-Aldrich Co.) to stock concentrations of 1 mg/mL and maintained at -70°C (max: 6 months).

### Minimum inhibitory concentration determination

The *in vitro* generated mutants were subjected to MIC testing using a limited range of 0.5, 1 and 2 µg/mL for both BDQ and CFZ using the BACTEC MGIT960 instrument, with results monitored on EpiCenter TBeXiST (Becton, Dickinson and Company, New Jersey, USA) as previously described (12, 13). When the GU of the drug-free GC reached 400 and if the GU of the drug-containing tube was ≥100, this was considered a resistant (R) result (12, 13). If the GU of the drug-containing tube was <100, the tube was incubated for a further seven days and if the GU of the drug-containing tube was ≥100 during this further 7 days of incubation (after the GU of the drug-free control tube reached 400), the strain was intermediate (I) as previously described (14), if it was still <100, the strain was susceptible (S) (12, 13).

### Creation of heteroresistant cultures

Mutant strains for which BDQ and CFZ resistance were confirmed by pDST, and their corresponding progenitors (wild type, wt) were grown in MGIT-OADC tubes (Figure 1) to the same time-to-positivity to ensure that both cultures entered the exponential growth phase when population mixtures were prepared. For each of the seven mutants a set of four heteroresistant cultures were prepared with a mutant:wt ratio mix of approximately 1:100, 5:100, 10:100, and 20:100 corresponding to a 1%, 5%, 10%, or 20% subpopulation of a BDQ-resistant clone.

### Phenotypic drug-susceptibility testing

For each of the heteroresistant cultures a corresponding 1:100 growth control (GC) was prepared in saline to represent 1% growth relative to the undiluted inocula. Five hundred microliters from this 1:100 dilution was transferred into the respective drug-free MGIT-OADC tubes. From each of the undiluted four heteroresistant mixtures 0.5 mL was transferred into two MGIT-OADC tubes containing 1.0 µg/mL or 2.0 µg/mL BDQ. For quality control undiluted wild-type or mutant strains were included to confirm their respective susceptibility and resistance when grown in BDQ-free and 1 µg/mL BDQ-containing MGIT-OADC tubes. A total of 28 assays were established (seven mutants at four different ratios) with a drug-free (GC) and two drug-containing tubes (test) for each assay (84 MGIT tubes in total, Figure 1).

### Determination of pDST profile and analysis of growth units

pDST results were determined for each heteroresistant mixture by using the GU and TTP data obtained by the BACTEC MGIT960 EpiCenter TBeXiST system as described above (12, 13). The time taken to reach 400 GU was also compared between all GC tubes for the various mixtures for each of the seven mutants and minimal deviations were seen between time-to-positivity for growth in drug-free GC tubes with differing ratios of mutant:wt (Table 1). Following determination of the pDST results, the entire volume from the 20% heteroresistant GC MGIT cultures (BDQ-free) were centrifuged and the pellet subjected to DNA extraction using the CTAB DNA extraction method as previously described (15, 16). The entire volume from the 10% heteroresistant GC MGIT cultures (BDQ-free) were centrifuged and the supernatant removed, except for 500 µL. This remaining volume was then subjected to thermal lysis at 100°C for 40-60 minutes.

### Pilot study for heteroresistant culture quantification

To determine whether heteroresistant mixtures could be reproducibly created and quantified with targeted NGS (tNGS), 5 ml of each of the heteroresistant mixtures (1%, 5%, 10%, or 20%) were created, and were split into two sets (2.5 mL) (Supplementary Figure S1). Set A was subjected to thermal lysis (1.25 mL) and pDST (1.25 mL) and set B was subjected to DNA extraction (1.25 mL) and pDST (1.25 mL). Thus, DST was performed in duplicate (set A and set B). Thermal lysates (set A) or pure DNA (set B) from heteroresistant mixtures prior to DST (1%, 5%, 10% and 20%) and the GC tubes post-DST (10% and 20%) were subjected to tNGS.

### Next generation sequencing of heteroresistant cultures

Both thermal lysates and pure DNA were shipped to the Translational Genomics Institute North in Arizona (USA) and used for targeted next generation sequencing of the *MmpR5* gene using a tiled, universal tailed method as previously described (17, 18). Data analysis was performed using the Amplicon Sequencing Analysis Pipeline (ASAP) with Single Molecule Overlapping Read (SMOR) technology as previously described (17, 18). Variants comprising at least 1% of each sample were reported using this software, which requires that forward and reverse sequencing reads agree to eliminate error. This ensures high confidence in variant calls. Variant frequencies were used to ensure that the dilutions created were in the expected range.

## Results

### Mutant characterization

Seven purified mutant strains with *MmpR5* variants were selected for further use, these included four mutants (Mutants 1, 2, 5, 6 and 7) derived from the ATCC25828 progenitor strain with *MmpR5* variants: A97G; G101T; 193delG; G287C and G326C and two mutant strains (Mutants 3 and 4) derived from ATCC27294 with *MmpR5* variants G126A and 192insC (Table 1). All seven were determined to be CFZ-resistant (MIC values of 2 µg/mL) and cross-resistant to BDQ (MIC values >2 µg/mL). Mutant 3 with a nonsynonymous variant (G126A: W42stop) and mutants 4 and 5 with frameshift mutations in *MmpR5* (192insC or 193delG) all possess WHO Group 2 variants (3). Mutants 1, 2, 6 and 7 have nonsynonymous *MmpR5* variants resulting in amino acid substitutions: A97G(T33A) and G101T(R24L) are ungraded variants (reported but not graded), G287C: R96P is a novel variant (not reported previously) and G326C: R109P is a WHO Group 3 variant (i.e. having an uncertain association with BDQ resistance) (3).

### Targeted next generation sequencing (tNGS)

Targeted next generation sequencing (tNGS) served as a confirmatory assay to quantify mutant:wt ratios, thereby defining the input material. A pilot study (Figure S1) comparing thermal lysates to DNA extracts showed that the 1% and 5% mixtures were too low for accurate detection by tNGS (Supplementary Table S1). DST results were comparable when performed in duplicate. For further tNGS assays, thermal lysates from the 10% GC tube and pure DNA from the 20% GC tube were chosen as inputs as these were determined to effectively quantify mutant:wt ratios. Analysis of the tNGS data showed that mutant *MmpR5* DNA was detected at an average of 17% (range, 10–30%) compared to the expected 20% from the drug-free GC (Table 1, Figure S2). Thermal lysates from drug-free GC tubes showed an average of 7% (range, 4–13%) of *MmpR5* mutant DNA compared to the expected 10% (Table 1, Figure S2).

### Phenotypic drug susceptibility testing of heteroresistant mixtures

Each mutant:wt mixture underwent four distinct pDST assays (with corresponding drug-free GC, 1 µg/mL and 2 µg/mL BDQ-containing MGIT tubes, Figure 1). The average time-to-positivity for the drug-free growth controls for all mixtures was 7.4 days (range (SD), 5.3–9.7 days (0.9 days)). At a 1 µg/mL concentration of BDQ, a resistant result was obtained from mutant 1 from the 1% mixture; from mutants 2, 4 and 6 from the 5% mixture, from mutant 5 from the 10% mixture and from mutants 3 and 7 from the 20% mixture (Figure 2). At lower ratios, mutants displayed an intermediate result (growth units in the drug-containing tube reached the threshold (≥100) for resistance but only after a further week of incubation) at 1 µg/mL of BDQ (Figure 2 and Supplementary Table S2). At a 2 µg/mL concentration of BDQ, resistant results were obtained for mutants 1, 2 and 6 from the 5% mixture, for mutant 3 from the 10% mixture and from mutants 4, 5 and 7 from the 20% mixture (Figure 2). At lower ratios, the mutants displayed an intermediate result at 2 µg/mL of BDQ, except for mutants 3 and 7 which were susceptible from a 1% mixture (Figure 2 and Supplementary Table S2).

**Figure 2:**
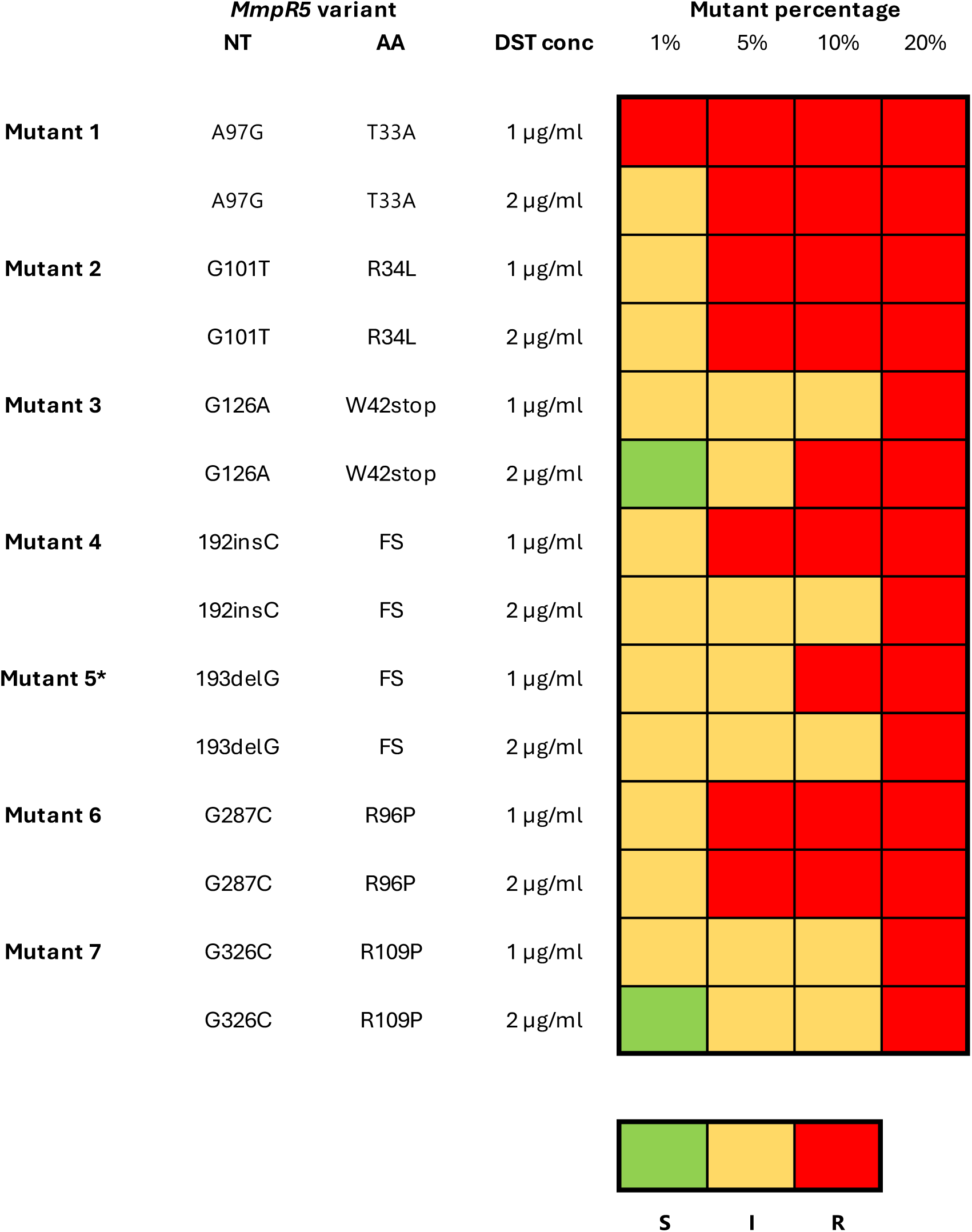
Heatmap showing the drug susceptibility profile at 1 and 2 µg/mL BDQ for each heteroresistant mixture at 1*, 5, 10 and 20% from 7 mutants. Susceptible pDST results are shown in green (S), intermediate pDST results in yellow (I) and resistant pDST results in red (R). Nucleotide changes (NT) and amino acid (AA) changes are shown for each mutant. Mutants 1 and 2 have *MmpR5* variants which have been previously described but are ungraded by the WHO, mutants 3-5 have *MmpR5* variants which have a group 2 WHO grading (associated with BDQ resistance in the interim), mutant 6 has a novel *MmpR5* variant and mutant 7 has a variant with a group 3 WHO grading, i.e. uncertain significance. *Mutant 5 DST result at 1% was determined using the GC from the 5% mutant as the TTP values were assumed to be similar and the GC for the 1% mixture failed due to contamination.

Figure 3 clearly highlights the importance of the intermediate category. The DST technique measures 99% inhibition of bacterial numbers to differentiate between resistance and susceptibility. Since resistant mutants were used for the preparation of the mixtures, the results show that at 2 µg/mL, we may fail to detect some of the resistant sub-populations in certain instances (e.g., mutants 3 and 7 which were susceptible from a 1% mixture). However, in most instances, using the current critical concentration (1 µg/mL) and the intermediate category, we can detect these resistant populations from as low as 1%. Furthermore, even without the use of the intermediate category, at 1 µg/mL we detected heteroresistant populations from as low as 5% in more than half the cases. Additionally, the intermediate results obtained for these mutants cross the threshold for resistance (GU ≥100) as early as 3 hours and up to 133 hours or 5.5 days after the corresponding GC has flagged positive (Supplementary Table S2).

**Figure 3:**
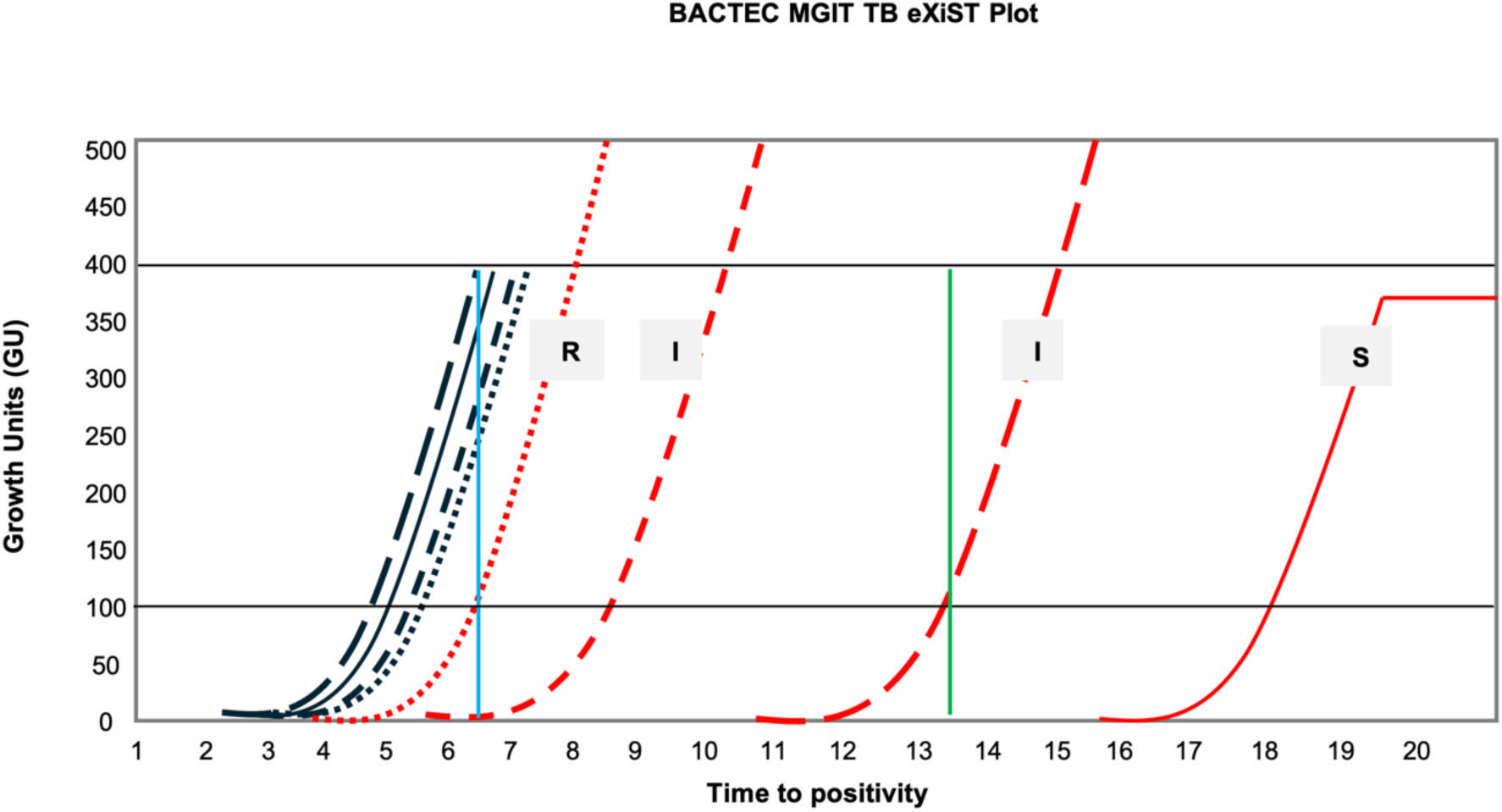
An example of a BACTEC MGIT960 EpiCenter TBeXiST plot for the experimental set-up used. Drug-free growth controls are shown in black for 1% (—), 5% (— —), 10% (- - -), and 20% (**^…^**) heteroresistant mixtures with corresponding BDQ-containing tubes (1 µg/mL BDQ) in red (with susceptibility result labeled). Minimal deviation is seen between growth controls, as the *MmpR5* variant has no impact in the absence of BDQ (i.e., the variant has no fitness cost). Horizontal black lines indicate either 100 growth units (GU; lower) or 400 GU (upper), the relevant thresholds as defined previously (12, 13). The vertical blue line distinguishes the exact separation point between R and I results, which is dependent on when the GC tube flags at 400GU (in this figure the 5% GC result is used, i.e. ∼6.5 days). The vertical green line indicates the 7-day period after which the growth control has flagged 400 GU (here, approximately day 14). Resistant (R), intermediate (I) or susceptible (S) results are reported when the corresponding GC has flagged positive at 400 GU and up to 7 days after. **R result:** the drug-free growth control tube reaches 400 GU and the drug-containing tube is ≥100 GU (12, 13). **I result:** the drug-free GC tube reaches 400 GU while the drug-containing tube only reaches ≥100 during the further seven days incubation (33). **S result:** the drug-free GC tube reaches 400 GU and the drug-containing tube remains <100 following 7 further days of incubation (12, 13). The plateau of the final exponential curve is indicative of the assay being ended in the instrument.

## Discussion

Following sixty years of use, pDST is still considered the gold standard for susceptibility testing and is currently implemented globally as a reflex standard-of-care assay. In this study, we show how the routinely used BACTEC MGIT960 platform coupled with EpiCenter TBeXiST software, can detect BDQ resistant subpopulations using available GU data. Through the extension of the incubation period of DST assays for a further seven days, intermediate results can be determined. Differences in the BDQ resistance profile for several mutants displaying a variety of *MmpR5* variants (including novel and Group 3 variants) were observed using the MGIT960 platform and two BDQ concentrations.

We attempted to explore the boundaries of the widely implemented MGIT960 assay to detect minor BDQ resistant subpopulations through the creation of low-level heteroresistant mixtures. While the MGIT960 assay is a qualitative assay exploiting fluorescence of an oxygen sensor for determination of inhibition of growth or lack thereof, we used several quantitative methods to validate our results and to ensure reproducibility for each of the seven mutants. This included dilution of a pure mutant culture to approximately 1, 5, 10 and 20%; tNGS from thermal lysates and pure DNA coupled with GU data and individual assays for each mutant at two different BDQ concentrations to maximise scientific rigor.

To simplify resistance classification, particularly in high-burden settings, a binary classification is used for MGIT960 results, i.e. either “S” for susceptible or “R” for resistant (19). Current binary classification methods categorize intermediate results as susceptible due to the 1% proportion rule, limiting the detection of minor resistant subpopulations. By extending the incubation period of MGIT960 DST assays by seven days, we were able to capture intermediate results indicative of heteroresistant populations. This can be achieved through interpretation of growth curve data (Figure 3), easily acquired from the BACTEC MGIT960 EpiCenter TBeXiST system which is universally used in conjunction with the MGIT960 (12, 13). It should be clarified that intermediate in this study is referring to the presence of <1% growth rather than a resistance category falling between susceptible and resistant categories. Therefore, heteroresistant populations should be easily detectable within the routine setting with data on susceptibility as well as any evidence of minor resistant populations within a week without drastic changes made to current testing algorithms. This method allows for more nuanced resistance profiling, offering a means to detect emerging resistance. Whether these resistant sub-populations are clinically significant for BDQ treatment does warrant further investigation, however, a study by Colangeli *et al* (20), showed that elevated isoniazid or rifampicin MIC values (below clinical breakpoints) were associated with greater risk of relapse in pretreatment isolates.

The current MGIT960 breakpoint for BDQ was established based on limited data and remains a matter of contention (7, 21), coupled with variants which have borderline MICs and the technical variability of phenotypic testing (7); BDQ DST faces multiple obstacles. To account for these challenges, we also made use of an additional concentration, 2 µg/ml BDQ, to enable a more granular interpretation of the resistance profiles for various heteroresistant mixtures. Although the phenotypic heterogeneity of underlying resistant populations is readily observed through growth curve data, the use of this higher concentration also shows that a limit of detection suitable to identify heteroresistance could be based on criteria that are different to those used for DSTs, i.e. a higher concentration could differentiate heteroresistant populations with greater clarity. Importantly, we used mutants with variants across the spectrum of the WHO grading criteria, i.e. LoF or frameshift variants, novel variants, reported but ungraded variants and variants with uncertain association with BDQ resistance (3). The use of the intermediate category for identification of heteroresistant populations proved to be valuable in all instances.

Several limitations exist when working with *M. tuberculosis* cultures. First, the bacteria are prone to clump in liquid MGIT culture with a “breadcrumb” appearance (22). We addressed this as best as possible with the creation of uniform mixtures through vortexing, allowing the mixtures to settle and using the supernatant. Second, due to the slow-growing nature of *M. tuberculosis*, a 24-hour difference in the time-to-positivity of the growth control tubes would represent a change equivalent to half of the bacterial population and this variability can be observed between GC tubes (Supplementary Figure S2). Finally, it appears that not all mutants return the same resistance profile; this could be due to differential growth in the mutant:wt mixture ratios (exhibited in differing tNGS percentages, (Supplementary Figure S2) or by the fact that not all *MmpR5* variants are equally responsible for high-level resistance (23). This may also be attributed to the degree by which specific *MmpR5* variants increase efflux pump activity and cause a reduction in the effective intracellular BDQ concentration. These possible scenarios were overcome using *in vitro* generated mutants, with *MmpR5* variants all associated with phenotypic resistance (i.e., an MIC of >1 µg/mL) as well as using mutant and progenitor cultures grown to the same log phase prior to heteroresistant mixtures were created.

Heteroresistance, characterized by mixed mutant and wild-type populations, is increasingly recognized as a challenge in TB treatment. In the case of BDQ heteroresistance, several studies have used next-generation sequencing to show the clinical impact of low-frequency variants in *MmpR5* (24–27). Previous studies have shown that in the absence of a supporting regimen to prevent resistance acquisition, *MmpR5* variants over time may lead to phenotypic resistance and poor treatment outcomes (28–31); an intermediate result would presumably have similar associations in this context. Ideally, an NGS technology utilized directly on clinical specimens and capable of detecting *MmpR5* and concurrent *MmpL5-S5* variants below 25% should be the reflex test for determination of BDQ resistance. If a variant with an uncertain association is identified, pDST should be repeated or MIC should be performed using an inoculum from the intermediate MGIT tube. This could result in an “R” pDST result, confirming the presence of heteroresistant populations (32) or elevated MIC values. The use of this composite reference standard would allow variants to be contributed to the WHO for the update of the mutation catalogue as well as improve time to detection of resistance. In summary, the ability to identify heteroresistant populations using existing diagnostic tools has important implications for TB resistance surveillance and treatment strategies. Future studies should focus on validating these findings with clinical samples and integrating them into routine DST workflows to improve early detection and patient outcomes.

## Acknowledgements

Fahd Naufal is acknowledged for administrative and project management support.

---

**Supplementary Table S1:** Pilot study (See Figure S1) time-to-positivity (TTP) data, as well as tNGS performed on thermal lysates (TL, Set A) or CTAB DNA extracts (DE, Set B). TTP refers to the time taken for the GC tube to reach 400 growth units (GU) was determined for each heteroresistant mixture. R result: the drug-free GC tube reaches 400 GU and the drug-containing tube is ≥100 GU. I result: the drug-free GC tube reaches 400 GU while the drug-containing tube only reaches ≥100 GU during the further seven days incubation (33). S: the drug-free GC tube reaches 400 GU and the drug-containing tube remains <100 GU following 7 further days of incubation. tNGS was used to determine the percentage of *MmpR5* detected in each heteroresistant mixture. ND: not detected, NA: not applicable.

| Thermal lysates | Time taken for GC to reach 400 GU (hours) | Drug susceptibility profile |  | % *MmpR5* variant detected with tNGS from thermal lysates (TL) |  |
| --- | --- | --- | --- | --- | --- |
| % | No Drug | 1 µg/mL BDQ | 2 µg/mL BDQ | Before DST | After DST from GC tube |
| 1% | 207 | I | S | ND | NA |
| 5% | 188 | I | I | 2.41 | NA |
| 10% | 206 | I | I | 6.58 | 10.41 |
| 20% | 244 | R | R | 12.29 | 20.83 |
| DNA extracts | Time taken for GC to reach 400 GU (hours) | Drug susceptibility profile |  | % *MmpR5* variant detected with tNGS from DNA extracts (DE) |  |
| % | No Drug | 1 µg/mL BDQ | 2 µg/mL BDQ | Before DST | After DST from GC tube |
| 1% | 189 | I | I | ND | NA |
| 5% | 205 | I | I | ND | NA |
| 10% | 204 | I | I | 2.36 | 2.52 |
| 20% | 237 | Contaminated | R | 12.36 | Contaminated |

**Supplementary Table S2:** Table showing growth units (GU) and time (days; hours) taken for growth control to reach 400 GU, as well as the GU and time (days; hours) recorded for drug-containing tubes when GC reached 100 GU and when drug-containing tubes reached 100 GUs for intermediate results only.

|  | Percentage mutant in mixture | Growth Control tube | 1 µg/mL BDQ-containing tube |  | 2 µg/mL BDQ-containing tube |  |
| --- | --- | --- | --- | --- | --- | --- |
|  |  | GU and time (days; hours) to reach 400 GU | GU and time (days; hours) recorded when GC reaches 400 GU | GU and time (days; hours) to reach 100 GU | GU and time (days; hours) recorded when GC reaches 400 GU | GU and time (days; hours) to reach 100 GU |
| Mutant 1 | 1% | 404 (7; 19) |  |  | 0 (7;19) | 107 (9;6) |
| Mutant 2 | 1% | 417 (6;20) | 76(6;20) | 101 (6;23) | 0 (6;20) | 100 (8;1) |
| Mutant 3 | 1% | 400 (7;5) | <12 (10;2) | 104 (10; 17) |  |  |
|  | 5% | 407 (7;15) | 53 (7;16) | 105 (7; 23) | <11 (12;10) | 101 (13; 4) |
|  | 10% | 404 (7;14) | >53 <106 (7;12 – 7;19) | 106 (7; 19) |  |  |
| Mutant 4 | 1% | 406 (7;14) | 0 (7;14) | 104 (8;12) | 0 (7;14) | 101 (12;9) |
|  | 5% | 420 (8;5) |  |  | <12 (8;19) | 100 (9;9) |
|  | 10% | 416 (7;3) |  |  | <10 (7;13) | 105 (8;5) |
| Mutant 5* | 1% | 414 (7;2) | <11 (8;6) | 100 (8;20) | <12 (9;11) | 106 (10;0) |
|  | 5% | 409 (7;0) | >10 <56 (6;21 – 7;7) | 108 (7;13) | <11 (7;12) | 107 (8;5) |
|  | 10% | 412 (6;21) |  |  | 53 (6;22) | 101 (7;5) |
| Mutant 6 | 1% | 423 (6;8) | 0 (6.8) | 101 (7;9) | 0 (6;8) | 101 (8;3) |
| Mutant 7<sup>a</sup> | 1% | 400 (8;15) | - | 100 (10;12) | - |  |
|  | 5% | 400 (7;20) | - | 100 (9;22) | - | 100 (13;0) |
|  | 10% | 400 (8;9) | - | 100 (8;21) | - | 100 (11;2) |

\*The control for the 1% mutant mix was contaminated so the GUs of the 5% mutant mix were used as a proxy.
<sup>a</sup>These results were taken from Graphs as Worklist GU values Trend Results are lacking.

## Supplementary methods

### PCR amplification

PCR amplification of the *M. tuberculosis Rv0678* gene was performed using forward (5’ AGAGTTCCAATCATCGCCCT 3’) and reverse primers (5’ TGCTCATCA GTC GTCCTCTC 3’).

Each 25 µL reaction solution comprised of nuclease-free water (8.5 µL), HotStarTaq® Plus Master mix (2X) (Qiagen, Hilden, Germany) (12.5 µL), 0.5 µl of each primer (10 pmol/ µL), 2 µL SYTO9 stain (ThermoFisher Scientific, Massachusetts, United States) and DNA (1 µL). A no-template control (NTC) and two positive controls consisting of pure and crude DNA from *M. tuberculosis* H37Rv were included in the assay. The amplification protocol consisted of an initial activation step of 95 °C for 5 minutes, followed by 40 cycles of 94 °C for 1 minute, 58 °C for 1 minute and 72 °C for 1 minute, followed by 72 °C for 10 minute and a melting step of 80°C for 15s and 95 °C for 15s with a change of 0.5° C/s increments was used to confirm amplification. All reactions were performed using QuantStudio 5 (Thermo Fisher Scientific).

### Phenotypic Drug Susceptibility Testing (pDST)

The *in vitro* generated mutants were subjected to MIC testing using a limited range of 0.5 – 2 µg/mL for both BDQ and CFZ. Five hundred microliters of standard inocula, prepared in MGIT-OADC tubes were used two days after the tubes flagged positive (400 growth units), was transferred to drug-containing MGIT-OADC tubes. A 1:100 (1%) dilution of the standard inocula was also prepared and 500 µL transferred to a drug-free MGIT-OADC tubes to serve as a growth control (GC). The tubes were then entered into the MGIT960 instrument, incubated at 37^°^C and results were subsequently monitored on EpiCenter TBeXiST (Becton, Dickinson and Company, New Jersey, USA). A susceptible *M. tuberculosis* H37Rv strain was included in each batch of BDQ and CFZ MIC determinations for quality control purposes. Tubes were incubated for at least 7 days after the 1:100 GC reached 400 growth units (GUs). Susceptibility of the cultures was determined and recorded according to the 1% proportion method as previously described (12) (Table 1).

### Next generation sequencing of heteroresistant cultures

Both thermal lysates and pure DNA were shipped to the Translational Genomics Institute North in Arizona (USA) and used for targeted next generation sequencing of the *Rv0678* gene using a tiled, universal tailed method as previously described (17, 18). Briefly, tailed primers (27) targeting *Rv0678* were used to amplify the full gene in a tiled approach. A second PCR step facilitated addition of a sequencing adapter via the universal tail. Libraries were pooled equimolarly and run on an Illumina NextSeq1000 using 2x 300bp, paired end chemistry, with a targeted coverage of 20,000 reads/amplicon. At least 30% of each sequencing run was filled with PhiX to ensure adequate base diversity for sequencing. Multiple no-template controls, as well as positive controls derived from H37Ra, were included with each run to ensure integrity of results. Data analysis was performed using the Amplicon Sequencing Analysis Pipeline (ASAP) with Single Molecule Overlapping Read (SMOR) technology as previously described (17, 18). Variants comprising at least 1% of each sample were reported using this software, which requires that forward and reverse sequencing reads agree to eliminate error. This ensures high confidence in variant calls. Variant frequencies were used to ensure that the dilutions created were in the expected range.

---

**Supplementary Figure S1:**
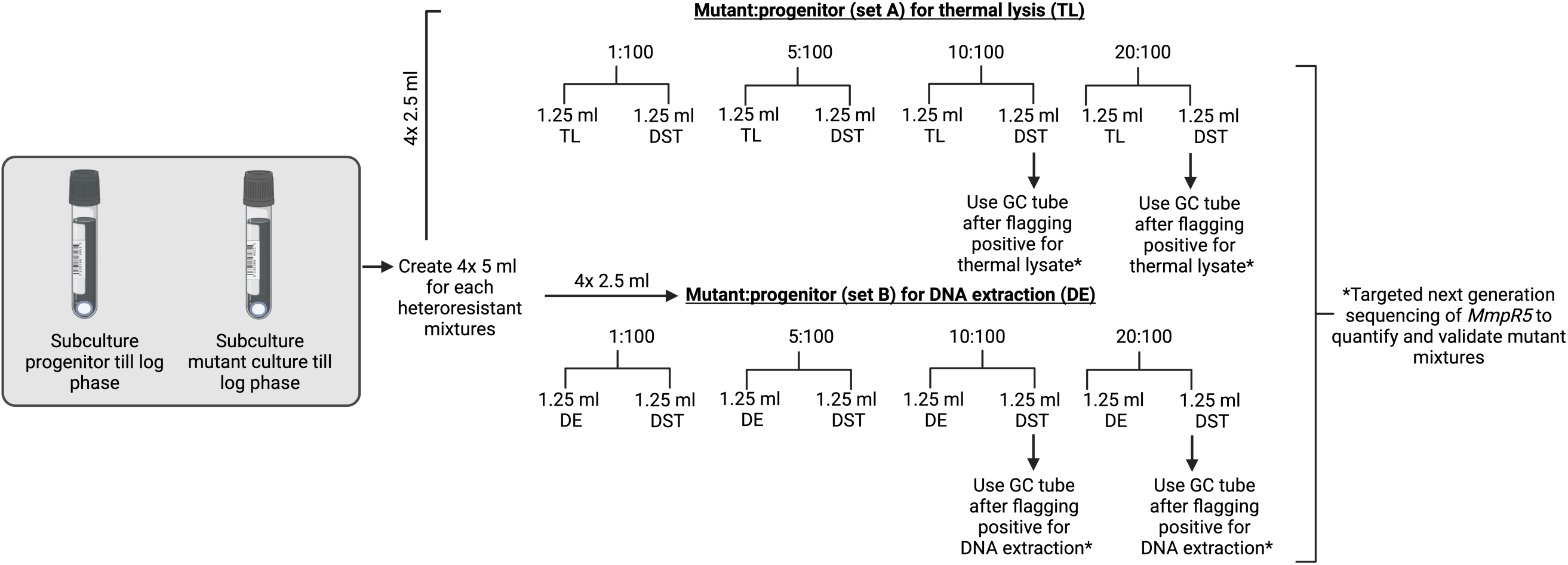
Pilot study to determine whether heteroresistant mixtures could be reproducibly created and quantified with targeted next generation sequencing. A single mutant was used for this experiment. 5 ml of each of the heteroresistant mixtures (1%, 5%, 10%, or 20%) was created, and split for either thermal lysis (set A: 1.25 ml, TL) or CTAB DNA extraction (set B: 1.25 ml, DE) (1.25 ml). The remaining 1.25 ml was then subjected to DST in duplicate (set A and set B). Thermal lysates and pure DNA were sent for tNGS of *MmpR5* to quantify mixtures. Created in https://BioRender.com

**Supplementary Figure S2:**
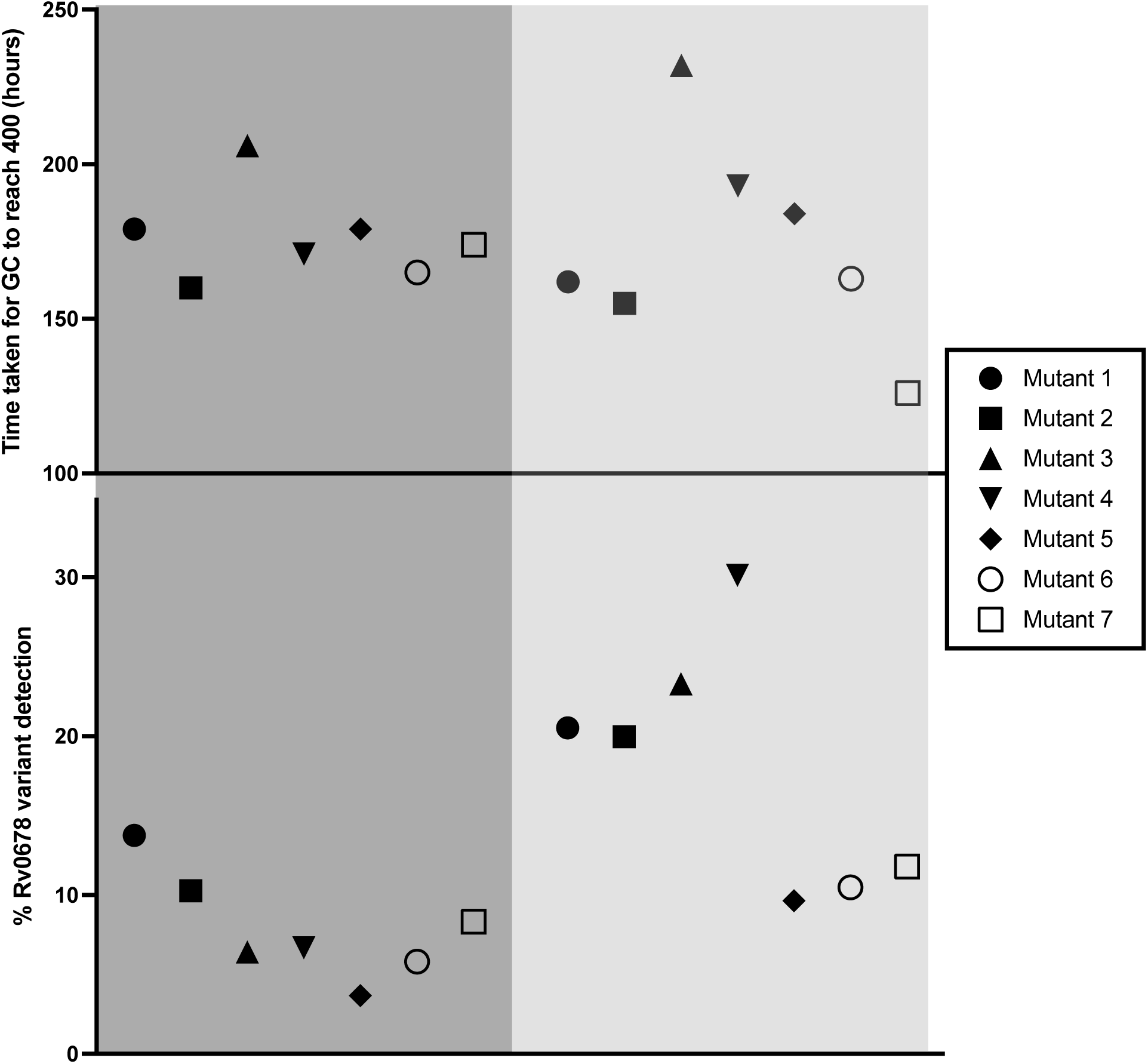
**Top:** Graph showing time taken for GC to reach 400 GU for 10% (dark grey) and 20% (light grey) heteroresistant mixtures. **Bottom:** Graph showing percentage MmpR5 detection from thermal lysates extracted from 10% heteroresistant mixtures (dark grey) and pure DNA extracted from 20% heteroresistant mixtures.

